# Medlib: A Feature-Rich C/C++ Library for Exact Alignment of Nanopore Sequences Using Multiple Edit Distance

**DOI:** 10.1101/2025.05.01.651420

**Authors:** Mehdi Rafeie, Fatemeh Vafaee, Omid R. Faridani

## Abstract

Pairwise sequence alignment is a cornerstone of bioinformatics, driving applications from variant calling to barcode demultiplexing. However, the advent of long-read, high-error sequencing platforms—such as Oxford Nanopore Technologies—demands alignment tools that not only deliver exact results but also accommodate the elevated noise inherent in these data. Here, we introduce Medlib (Multiple Edit Distance Library) that offers both high performance and novel, biologically motivated features. Medlib’s threshold mode allows users to retrieve every alignment with edit distance k ≤ k_th_ and optionally alignment score s ≥ s_th_, ensuring comprehensive discovery of all plausible matches. Counterintuitively, this exhaustive retrieval outpaces traditional minimum-error searches. Medlib further provides the first exact implementations of overlap-aware reporting and motif boundary treatment. Medlib supports multiple input formats, predefined and custom alphabets, affine-gap scoring applied retrospectively, built-in dedicated multithreading schemes, and user-defined function callbacks for immediate post-processing. Benchmarking against Edlib demonstrates a 1.6× speedup in minimum mode and even greater gains (2.5×) in threshold mode—while delivering up to twice as many alignments. Medlib thus offers a robust, feature-rich engine for exact pairwise alignment in modern, high-throughput sequencing workflows.

**Availability and Implementation:** Source code, installation instructions and test data are freely available for download at https://github.com/Mehdi-Rafeie/Medlib, under the MIT license.

## Introduction

Pairwise sequence alignment is a foundational operation in bioinformatics, underpinning critical analyses such as variant calling [1], barcode demultiplexing [2], transcriptome reconstruction [3], genome assembly [4], homology detection [5], and comparative genomics [6]. Accurate alignments are essential for downstream tasks including structural-variation discovery and single-nucleotide polymorphism identification, where even small errors can propagate through complex analytical pipelines. As sequencing technologies have advanced, researchers have shifted from short-read platforms to third-generation approaches that provide much longer reads but also exhibit higher base-calling error rates [7].

Nanopore-based sequencing, in particular, is a transformative technology that enables direct, real-time analysis of single DNA [8] or RNA molecules as they pass through nanoscale pores. Unlike traditional short-read platforms, it can produce ultra-long reads—often spanning hundreds of kilobases—making it particularly valuable for resolving complex genomic regions, structural variants, and full-length transcripts. A key advantage of nanopore sequencing is its ability to detect native DNA and RNA modifications such as methylation [9, 10], which are not accessible with other sequencing technologies. Beyond nucleic acids, nanopore applications are rapidly expanding into proteomics, with recent advances demonstrating its potential for single-molecule protein sequencing and detection of post-translational modifications [11]. Despite these strengths, nanopore sequencing still faces a major limitation: a relatively high error rate compared to other platforms, often exceeding 10% [12, 13]. These errors often stem from systematic signal biases and stochastic noise, which can compromise the accuracy of variant calling. While ongoing improvements in pore chemistry, sensor design, and base-calling algorithms continue to reduce error rates, the remaining noise level poses significant challenges for alignment algorithms that were originally designed for low-error short reads.

Over the past four decades, exact alignment algorithms have progressed from the Levenshtein distance formulation—which counts the minimal number of insertions, deletions or substitutions required to transform one sequence into another [14]—to global and local dynamic-programming (DP) methods. The Needleman–Wunsch algorithm achieves optimal global alignment [15], and the Smith–Waterman algorithm performs optimal local alignment [16], both were extended by Gotoh to support affine gap penalties, providing a more biologically realistic model of insertion and deletion costs [17]. However, these DP approaches have time and space complexity proportional to the product of the sequence lengths, O(n·m), which becomes prohibitive for ultra-long reads [18]. To mitigate this, algorithmic optimizations such as wavefront alignment (WFA) restrict computation to promising matrix regions [18], while modern CPU architectures exploit SIMD vectorization and multithreading [19, 20], and GPUs offer hardware acceleration [21, 22]. Complementary innovations include BSAlign, a recent ultra-fast library that exemplifies the synergistic application of striped SIMD vectorization, the difference-recurrence relation, and banded DP to achieve substantial speed gains [23].

In parallel, heuristic methods—fixed-width banding, seed-and-extend schemes (e.g., BLAST [24], BWA [25]) and drop-off heuristics like X-drop [26] and Z-drop [27]—have been widely adopted for rapid, approximate alignment. These strategies can miss true alignments when indels shift optimal paths outside predefined bands or when extension scores fall below arbitrary thresholds. An exception to this trade-off is QuickEd, a recent aligner representing a hybrid approach: it leverages fast heuristic bounding to achieve record-breaking throughput before computing the exact optimal alignment, but ultimately reports only the single best match [28].

Most exact aligners continue to prioritize the minimum-error alignment, discarding all other near-optimal candidates. While efficient, this best-match paradigm can overlook biologically meaningful alternatives—especially in high-error contexts such as noisy long reads, barcode identification, or conserved motif discovery—where multiple alignments within an acceptable error margin may be relevant. Addressing this gap requires a tool that combines exactness with inclusive retrieval.

Here, we present Medlib, a high-performance C/C++ library for exact pairwise sequence alignment that builds directly on the foundation of Edlib [29], extending Myers’s bit-vector algorithm [30] with novel threshold-based retrieval, overlap awareness, and motif boundary sensitivity. Medlib’s Threshold operation mode enables users to retrieve every alignment with edit distance *k* ≤ *k*_th_ (where *k*_th_ is a user-defined maximum edit distance) and optionally alignment score *s* ≥ *s*_th_ (where *s*_th_ is a user-defined minimum alignment score), ensuring comprehensive discovery of all plausible matches without resorting to heuristics.

Counterintuitively, this exhaustive mode can outperform traditional minimum-error retrieval, as it omits the post-filtering phase required to isolate the best match. Beyond thresholding, Medlib offers the first exact implementation of overlap-aware reporting, distinguishing redundant from distinct alignments, and flexible motif boundary treatment, which reinterprets mismatches at query ends as deletions to preserve the neighboring base pairs for an adjacent barcode.

Medlib’s architecture is built on a policy-based design that parameterizes core alignment logic at compile time, eliminating runtime conditionals and supporting static polymorphism via template metaprogramming. A compile-time dispatch mechanism instantiates only the required code paths, and a selective compilation interface mitigates build-time explosion when hundreds of alignment modules are available. The library natively supports multiple input modalities (in-memory sequences, plain-text, FASTA and FASTQ), predefined and user-defined alphabets, affine-gap scoring applied retrospectively to Myers-derived edit paths, and user-defined function callbacks for real-time post-processing. Dedicated multithreading schemes tailored to each alignment type maximize throughput on multi-core systems, and comprehensive status codes paired with automatic configuration validation ensure robust, debuggable execution.

Together, these innovations establish Medlib as a versatile, exact alignment engine optimized for the unique challenges of noisy, long-read sequencing data and modern high-throughput bioinformatics pipelines. In the Method section, we describe Medlib’s design principles, feature set and performance benchmarks against widely used aligners.

## Method

### Policy-Based Design Pattern

Medlib adopts a policy-based design pattern to enable high configurability without sacrificing performance. Core alignment logic is parameterized through template policies that define behavior such as input types, operation modes, and threading strategies. These policies are instantiated at compile time based on user configuration, allowing customization while avoiding runtime conditionals. This design permits a modular yet efficient structure, simplifying extension and maintenance.

### Static Polymorphism

To minimize runtime overhead, Medlib replaces classical runtime polymorphism with static polymorphism through template metaprogramming. By resolving function calls and specializations at compile time, Medlib eliminates the cost of virtual dispatch and supports inline expansion and compiler optimizations. This results in a zero-overhead abstraction model that is particularly beneficial in performance-critical contexts such as high-throughput sequence alignment.

### Compile-Time Dispatch

Medlib centralizes control flow through a compile-time dispatch mechanism. Upon user configuration, a dispatcher instantiates the appropriate alignment function with the correct template arguments, ensuring only relevant code paths are generated and compiled. This approach ensures tight coupling between configuration and execution logic while enabling the compiler to apply aggressive optimizations.

### Selective Compilation

To accommodate the large number of supported functions—over 150 functions that expand to 850,000+ instantiations when compiled in full—Medlib provides a selective compilation interface. This mechanism allows developers to compile only the modules required for their specific use cases, controlled via the medlib_compile.h interface. By limiting the compilation scope, this feature mitigates the combinatorial explosion in build time and memory usage, especially in Debug mode. Full compilation (e.g., in Release mode) remains available for benchmarking or general-purpose deployments.

### Supported Input modalities and Alignment Types

Medlib offers extensive flexibility in how input sequences are provided and aligned, supporting a variety of query and target formats suited to diverse analysis pipelines. Queries can be specified as single sequences in memory, batches of sequences in memory, or plain-text files in which each line contains one query sequence. Targets can likewise be specified as single or multiple sequences in memory, or as sequence files in one of three supported formats: plain text (with each line containing a target ID followed by a comma and the sequence), FASTA, or FASTQ. These input modalities yield 15 distinct alignment types, representing all valid pairwise combinations of query and target formats. Each alignment type is implemented as a separate module, allowing for independent compilation and execution. This modular design enables fine-grained control over memory usage and runtime behavior, facilitating efficient integration into high-throughput or customized processing pipelines. For detailed descriptions of the alignment types, please refer to Table S1 in the Supplementary Information document.

### Supported Alphabets and Customization

In contrast to Edlib—which infers the alphabet dynamically at runtime—Medlib enables users to specify predefined alphabets explicitly. This design facilitates compile-time optimization and ensures compatibility with custom encoding schemes, particularly advantageous in specialized contexts such as barcode demultiplexing or proteomics. As in Edlib, Medlib also supports the definition of character equalities, allowing users to treat ambiguous or degenerate symbols (e.g., interpreting ‘N’ as matching any of A, C, G, or T) as equivalent during alignment.

### Threshold Operation Mode

In addition to the classic minimum edit distance retrieval mode—used by Edlib to identify only the closest alignment for each query—Medlib introduces a novel threshold-based operation mode. This mode retrieves all alignments whose k is less than or equal to a user-defined threshold, kThr. By shifting from a strict minimum search to a bounded range query, the threshold mode enables more inclusive alignment discovery, which is critical in applications where multiple near-optimal alignments are biologically meaningful (e.g., detecting sequence variants [ref], handling ambiguous barcodes [ref], or identifying conserved motifs [ref]).

The motivation for this mode is especially evident in noisy sequencing platforms such as ONT, where base-level error rates can exceed 10%. In such settings, one is often interested in identifying all alignments with an error rate below or equal to a specific threshold—such as 10% of the query length—rather than just the single best match. This inclusive approach is more robust to stochastic sequencing errors and improves sensitivity when downstream analyses benefit from capturing all plausible matches.

Internally, this mode retains the speed and accuracy of Myers’s bit-vector algorithm and leverages Ukkonen’s banding strategy to confine computation to the minimum necessary diagonal band. Unlike heuristic implementations of banded alignment, Medlib’s threshold mode guarantees completeness, ensuring that all qualifying alignments are discovered without compromising correctness.

### Overlap Handling and Distinct Alignments

In high-throughput sequence alignment tasks—particularly those involving short queries like barcodes or motifs—multiple alignments may overlap along the target sequence. Traditional alignment tools, including Edlib, do not distinguish between overlapping alignments, often tending to report all matches regardless of their positional redundancy. While this behavior aims to maximize sensitivity, it can obscure the biological interpretation of true alignment signals, inflate result counts with redundant matches, and complicate downstream processing. Moreover, due to the lack of overlap-aware logic, they may inadvertently miss valid alignment instances when nearby suboptimal matches overshadow true signals.

To address this, Medlib introduces an overlap-aware alignment framework that supports both inclusive and exclusive reporting of overlapping alignments. Specifically, Medlib allows users to choose between reporting All valid alignments or only those that are Distinct. A distinct alignment is defined as one that is separated from other reported alignments by at least kThr bases at either the start or end position. This ensures that each distinct result represents a unique, non-redundant alignment event.

This feature is especially useful in contexts such as barcode demultiplexing, motif discovery, or peak calling, where reporting multiple, highly similar overlapping hits can lead to ambiguity, overcounting, or bias in biological interpretation. By offering precise control over overlap resolution, Medlib enables users to tailor alignment reporting to the biological or analytical requirements of their pipeline.

Medlib’s overlap handling is fully integrated into both minimum and threshold operation modes, allowing users to maintain consistency and clarity in their alignment outputs regardless of the underlying search strategy.

### Motif Boundary Treatment

Accurate interpretation of motif alignments often requires special handling of mismatches occurring at the boundaries of the aligned regions. Traditional alignment tools typically treat all mismatches equally, without distinguishing whether they occur internally or at the start or end of the query motif. However, in many biological contexts— such as locating motifs flanked by variable sequences—mismatches at the motif boundaries are often tolerated because they can result from imprecise alignment against adjacent regions rather than true mutations within the motif itself. In contrast, internal mismatches are more likely to represent genuine biological variations and are treated accordingly.

To address this, Medlib introduces a flexible motif boundary treatment system that allows users to control how boundary mismatches are interpreted during alignment scoring. Four boundary treatment modes are supported: No Flanking, Flanking Start, Flanking End, and Flanking Both. In No Flanking mode, mismatches at the motif boundaries are treated normally, contributing to the k value like any internal mismatch. In Flanking Start/End modes, mismatches at the beginning/end of the motif are not penalized as mismatches but are reinterpreted as deletions from the adjacent upstream sequence, thereby preserving the motif’s integrity. In Flanking Both mode, this reinterpretation is applied at both the start and end boundaries simultaneously, ensuring that neither end of the motif absorbs bases that properly belong to neighboring sequence elements.

This boundary reinterpretation is particularly important in high-throughput omics workflows, such as single-cell assays relying on engineered constant regions (e.g., primers or linkers) to locate adjacent barcodes or UMIs. When aligning constant regions, aggressive ownership of boundary mismatches can cause bases from adjacent barcodes to be mistakenly absorbed into the aligned region, leading to missed or distorted barcode identification. By reinterpreting boundary mismatches as deletions from adjacent sequences, Medlib ensures that constant regions are delineated precisely, safeguarding the accurate detection of neighboring barcodes and preserving biological fidelity.

Motif boundary treatment is fully integrated into both Minimum and Threshold operation modes, allowing users to apply boundary sensitivity consistently across different alignment strategies. By supporting flexible, biologically meaningful handling of boundary mismatches, Medlib provides researchers with greater precision and robustness in motif- and barcode-centric analyses.

### Scoring Scheme and Affine Gap Penalties

While Medlib is fundamentally rooted in exact edit distance computation, it further extends its utility by supporting customizable scoring schemes, including affine gap penalties. In contrast to the basic Levenshtein model, biologically realistic alignments often require distinguishing between gap openings and gap extensions, reflecting the greater likelihood of contiguous indel events compared to isolated ones. Medlib incorporates this flexibility by optionally allowing users to define separate penalties for matches, mismatches, gap openings, and gap extensions, adopting a widely-used scoring model consistent with classical alignment algorithms such as Gotoh’s algorithm [17].

The scoring functionality is implemented as a lightweight, optional module layered over the primary alignment computation. After Medlib identifies the full edit path using its Myers-based engine, it retrospectively traverses the path to assign a cumulative alignment score according to the user-specified penalties. This post-alignment scoring step preserves the speed advantages of bit-parallel edit distance calculation while enabling users to rank or filter alignments based on biologically meaningful scoring models. By bridging the simplicity of edit distance calculations with the expressiveness of affine scoring systems, Medlib ensures that users can flexibly adapt alignment criteria to the needs of complex biological datasets without sacrificing computational efficiency.

### User-Defined Functions (UDFs)

Medlib enhances workflow flexibility by supporting UDFs, allowing developers to directly integrate custom post-processing logic into the alignment pipeline. Instead of relying solely on intermediate outputs such as result files—which can introduce significant I/O overhead—users can attach UDF callbacks that operate immediately on each alignment result as it is produced. The UDF system is fully type-safe and integrated at the template level, ensuring that no runtime penalty is incurred when user callbacks are not supplied. Developers retain full control over the structure and content of UDFs, enabling customized operations such as result parsing, scoring adjustments, metadata tagging, or direct insertion into downstream pipelines like variant annotation, barcode mapping, or clustering analyses. By providing immediate, programmable access to alignment outputs without redundant disk I/O or intermediate parsing, Medlib’s UDF framework bridges the gap between core alignment computation and customized downstream interpretation, fostering seamless integration into diverse bioinformatics workflows.

### Dedicated Multi-Threading for Each Alignment Type

To maximize throughput on large datasets, Medlib implements dedicated multi-threading strategies tailored to each alignment type. Medlib statically assigns optimized parallelization schemes at the function level, depending on the specific combination of query and target modalities. By tightly coupling the threading design to the data access patterns of each alignment type, Medlib minimizes synchronization overhead, maximizes CPU core utilization, and achieves near-linear scaling on multi-core processors. For detailed descriptions of the multi-threading strategies applied to each alignment type, please refer to Table S2 in the Supplementary Information document.

### Status Codes and Configuration Validation

Robust error handling and configuration validation are central to Medlib’s design philosophy. To support transparent diagnostics and facilitate debugging, Medlib defines a comprehensive set of status codes that report the outcome of operations ranging from configuration parsing to sequence alignment execution. Status codes are organized into distinct categories, including general operational statuses, configuration validation statuses, UDF statuses, and file operation statuses.

Configuration validation is performed automatically when a configuration object is received. Medlib systematically checks all user-provided parameters against an internal set of logical consistency rules, encompassing input types, output formats, parallelism settings, scoring configurations, motif treatment policies, and other operation-specific constraints. If any inconsistency or violation is detected, an appropriate status code is immediately returned, along with a descriptive error message. These validation steps ensure that misconfigured runs are intercepted early, preventing undefined behavior and saving computational resources during alignment.

Medlib also offers dedicated utility functions for retrieving human-readable descriptions of each status code, allowing developers and users to diagnose and correct issues efficiently without manually inspecting source code or logs. A detailed tree structure summarizing all configuration validation rules, along with their associated rule identifiers, is available at https://github.com/Mehdi-Rafeie/Medlib/ConfigRules.

### Experimental Design

We compare Medlib against two widely used alignment tools: Edlib (v3.6) [29] (for exact edit distance) and Parasail (v.2.4.3) [19] (for affine scoring). Additionally, we benchmark Medlib against BSAlign [23], previously reported as the fastest library among Myers [30], SSW [20], ksw2 (being a component of Minimap2 [27]), WFA [18], BA [31], as well as Edlib and Parasail. To ensure comparability across all applicable tools, we employed a unified scalar scoring system. Specifically, match, mismatch, gap open, and gap extension scores were consistently set at +2, −4, −4, and −2, respectively, in all experiments.

For consistency with BSAlign’s published benchmarks, we utilized the same real sequencing datasets. The short-read dataset comprises 100,000 pairs of 101-bp Illumina HiSeq 2000 reads (Experiment Accession: ERX069505). The long-read dataset consists of 12,477 Oxford Nanopore MinION reads, each approximately 1,000 bp in length (Experiment Accession: ERX3305687).

## Results

We conduct all benchmarks on a Windows 11 64-bit system equipped with an Intel Core i9-14900HX processor and 64 GB of random-access memory (RAM). All programs are compiled for 64-bit x86-64 architecture with AVX2 SIMD instruction set support. Execution time is measured as elapsed wall-clock time using C++ standard timing utilities in single-threaded mode. Each alignment experiment is repeated 1,000 times to ensure statistical stability. The recall rate is defined as the proportion of alignments that yield the same score as a trusted baseline tool—Edlib for edit distance mode, and Parasail for affine scoring mode—under an identical scoring scheme. We also record the maximum memory usage reported by the system during each benchmark. Overall, Medlib demonstrates competitive or superior performance compared to Edlib and other tools under the tested conditions.

### Performance Comparison with Edlib

We compare Medlib with Edlib using a large-scale alignment task involving a single query sequence aligned against 3,469,149 target sequences. The benchmarks are run in single-threaded mode without applying a scoring scheme, and execution time is measured as wall-clock duration using C++ standard timing utilities. To ensure consistency, each test is repeated 1,000 times.

In terms of execution time, Medlib outperforms Edlib across all evaluated operation modes. Specifically, in the Minimum mode—which is functionally equivalent to Edlib’s default behavior—Medlib completes the task in 1 minute and 12 seconds, compared to Edlib’s 1 minute and 22 seconds. This represents a 1.64× speedup. Notably, Medlib also produces a substantially higher number of alignments (21,623,281 vs. 15,012,036), suggesting that Edlib may miss valid results even in its exact mode. This discrepancy highlights Medlib’s enhanced ability to capture all qualifying alignments, potentially due to more comprehensive internal banding and path-retrieval logic.

Medlib’s performance advantage becomes even more pronounced in Threshold mode, which retrieves all alignments with edit distance ≤ k, instead of just the minimum-distance match. Despite its broader search space and increased result count (32,924,809 alignments), Medlib executes in just 1 minute and 7 seconds—yielding a 2.34× speedup compared to Edlib’s baseline. This highlights Medlib’s ability to maintain high performance even when performing substantially more work.

To the best of our knowledge, no existing alignment tool—including Edlib, Parasail, BSAlign, or WFA—implements an exact threshold-based mode that guarantees retrieval of all alignments up to a user-defined error threshold. Existing heuristics such as fixed-width banding, X-drop, and seed-extension methods offer approximate retrieval but do not ensure completeness. Thus, Medlib introduces a unique capability in the space of exact alignment tools.

These performance trends are summarized in Figure 1A, which shows the speedup of Medlib relative to Edlib across modes, and in Figure 1B, which illustrates the increasing alignment counts corresponding to Edlib, Minimum, and Threshold modes. Notably, while Medlib’s Threshold mode yields more than twice the alignments compared to Edlib, it still executes faster, demonstrating its scalability and efficiency.

**Figure 1.**
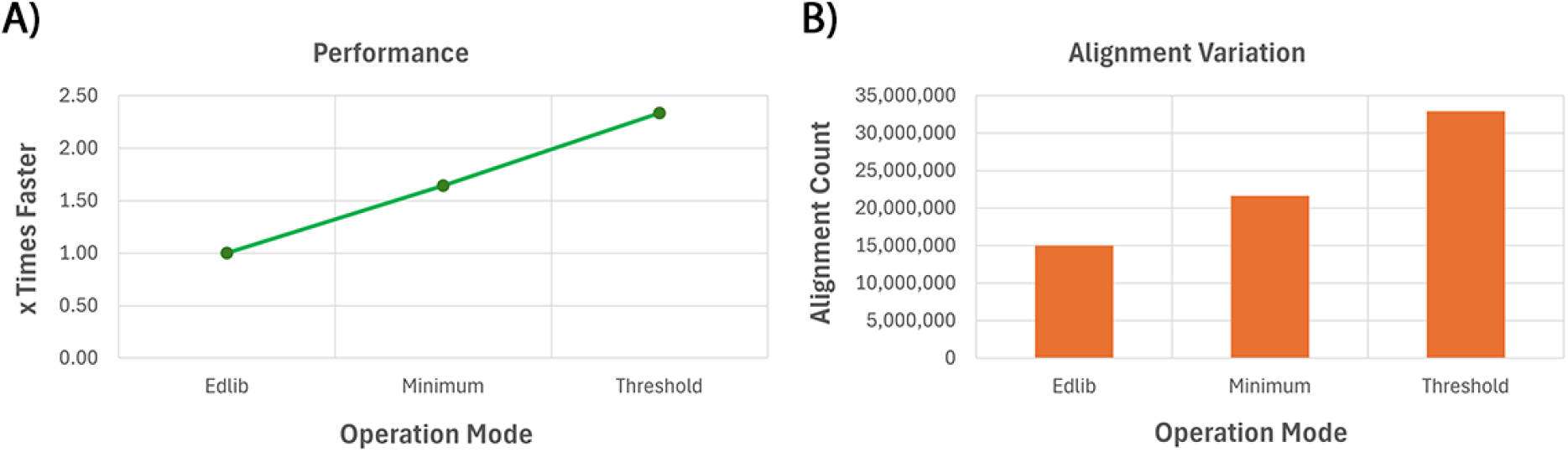
Comparative performance and alignment count across different operation modes. (A) Relative execution speed of Medlib in Minimum and Threshold modes compared to Edlib, expressed as fold-increase in performance. Despite performing broader searches, Medlib maintains superior speed, achieving up to 2.34× faster execution in Threshold mode. (B) Total alignment counts returned under each operation mode. Medlib yields substantially more alignments than Edlib in both Minimum and Threshold modes, indicating improved sensitivity. The Threshold mode, uniquely supported by Medlib, captures all alignments within a user-defined edit distance, resulting in over twice as many hits as Edlib.

### Overlap Handling Modes

To assess the impact of overlap resolution on performance and output variation, we benchmark Medlib in both All and Distinct modes and compare them to Edlib, which does not natively support overlap-aware filtering. In All mode, Medlib reports all alignments that satisfy the minimum distance criterion, including overlapping hits. In contrast, Distinct mode restricts output to non-redundant alignments, defined as results separated by at least k bases at the start or end position, thereby reducing ambiguity and redundancy in the output.

Compared to Edlib, which identifies 15,012,036 alignments, Medlib’s All mode reports 21,623,281 hits—representing a 44% increase—while completing the task in 1 minute and 12 seconds, yielding a 1.64× speedup. This demonstrates that Medlib can retrieve substantially more results with higher recall while maintaining superior performance.

When run in Distinct mode, Medlib filters overlapping alignments and returns 12,629,671 unique hits. While this operation incurs a slight runtime overhead (execution time increases to 1 minute and 15 seconds), the performance still remains close to Edlib, at 0.95× relative speed. Importantly, this mode offers a more conservative, biologically meaningful view of alignment results by eliminating locally redundant matches.

Together, these results demonstrate that Medlib not only improves alignment recall and execution speed in All mode, but also provides a tunable mechanism for users to reduce redundancy when clarity and interpretability are preferred. Figure 2A illustrates the performance trends, while Figure 2B visualizes the corresponding variation in alignment count across modes.

**Figure 2.**
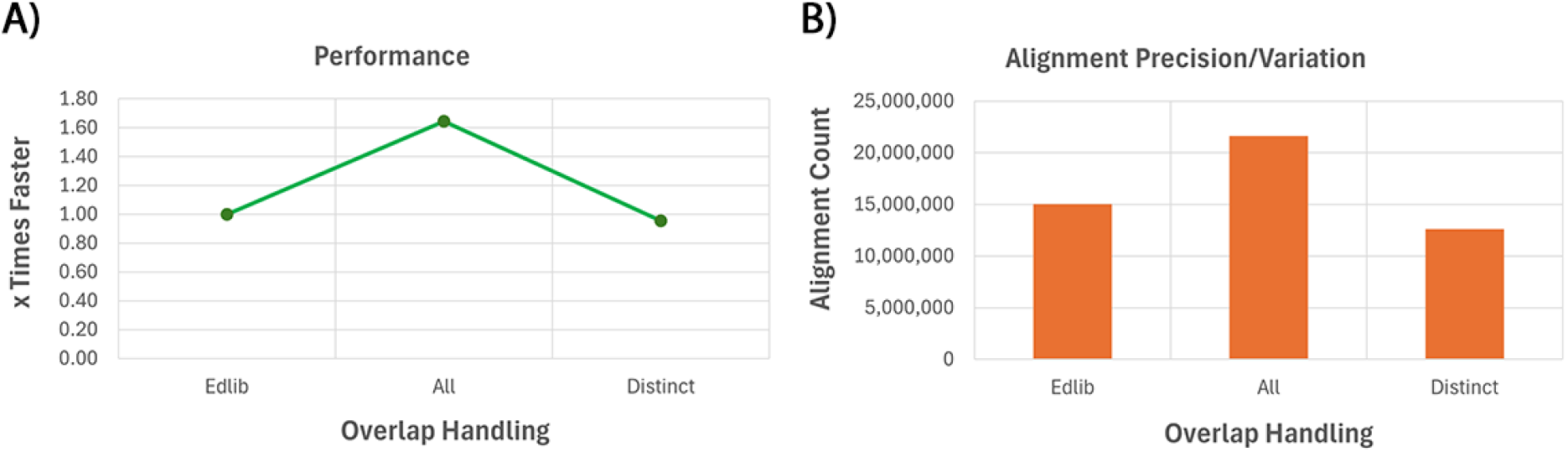
Impact of overlap handling on performance and alignment count. (A) Relative speed of Medlib in All and Distinct overlap handling modes compared to Edlib. Medlib maintains higher performance even when reporting all possible alignments. (B) Total number of alignments returned under each mode. Medlib’s All mode retrieves substantially more alignments than Edlib, while the Distinct mode produces fewer, non-overlapping alignments with minimal performance trade-off.

### Motif Boundary Sensitivity

To assess the utility and performance trade-offs of Medlib’s motif boundary treatment feature, we benchmark its four supported modes: No Flanking, Flanking Start, Flanking End, and Flanking Both. These modes allow users to selectively exclude mismatches occurring at the motif boundaries from penalization—mismatches that often arise due to imprecise read boundaries or adjacent variable regions.

As shown in Figure 3A, motif boundary treatment introduces moderate runtime overhead, particularly when applied symmetrically. The No Flanking mode, which penalizes all mismatches equally, completes execution in 1 minute and 13 seconds and serves as the baseline for comparison. Flanking Start and Flanking End modes increase runtime to 1:16 and 1:14, respectively, while Flanking Both takes 1:18. These timings correspond to relative performance slowdowns of 32% (Flanking Start), 21% (Flanking End), and 45% (Flanking Both) compared to the baseline.

**Figure 3.**
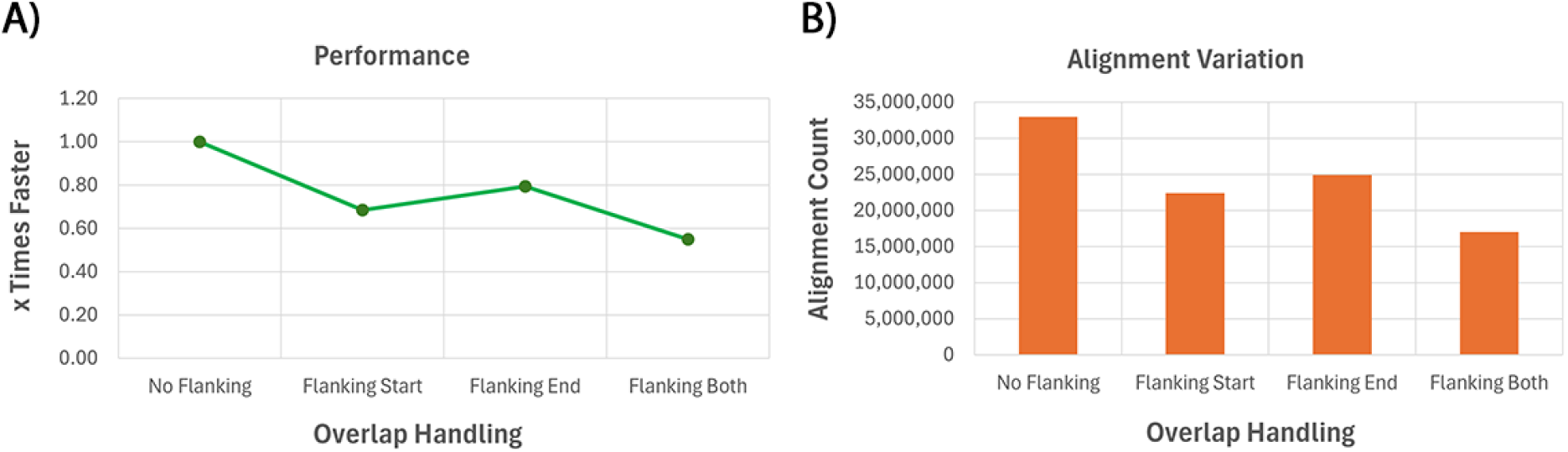
Runtime and alignment count under different motif boundary treatment modes in Medlib. (A) Relative execution time of each boundary mode, normalized to the No Flanking baseline. Treating boundary mismatches as deletions from adjacent sequences increases computational cost modestly. (B) Total alignment count under each mode. More permissive modes such as No Flanking yield higher hit counts, while stricter modes like Flanking Both reduce potentially spurious alignments at motif edges.

Functionally, boundary treatment has a strong impact on alignment count. As shown in Figure 3B, No Flanking yields the largest number of alignments (32,924,809), while the strictest mode, Flanking Both, yields the fewest (17,002,216). This substantial drop reflects Medlib’s ability to eliminate boundary-associated spurious alignments that would otherwise be counted as legitimate hits. Flanking Start and Flanking End offer intermediate levels of precision, producing 22,395,883 and 24,869,088 alignments, respectively.

Together, these results illustrate the value of motif boundary treatment as a biologically meaningful filter for refining alignments in applications such as barcode detection, motif discovery, or sequence annotation. Medlib’s implementation enables researchers to tailor alignment strictness with minimal performance cost while gaining substantial control over result specificity.

### Multi-Threading Efficiency

To assess the scalability of Medlib’s parallel execution model, we benchmark performance under increasing thread counts (1 to 4 threads) using the same input—a single query aligned against 18,000 target sequences—under identical scoring and filtering parameters. As shown in Figure 4, Medlib demonstrates consistent, near-linear improvements in execution time with additional threads.

**Figure 4.**
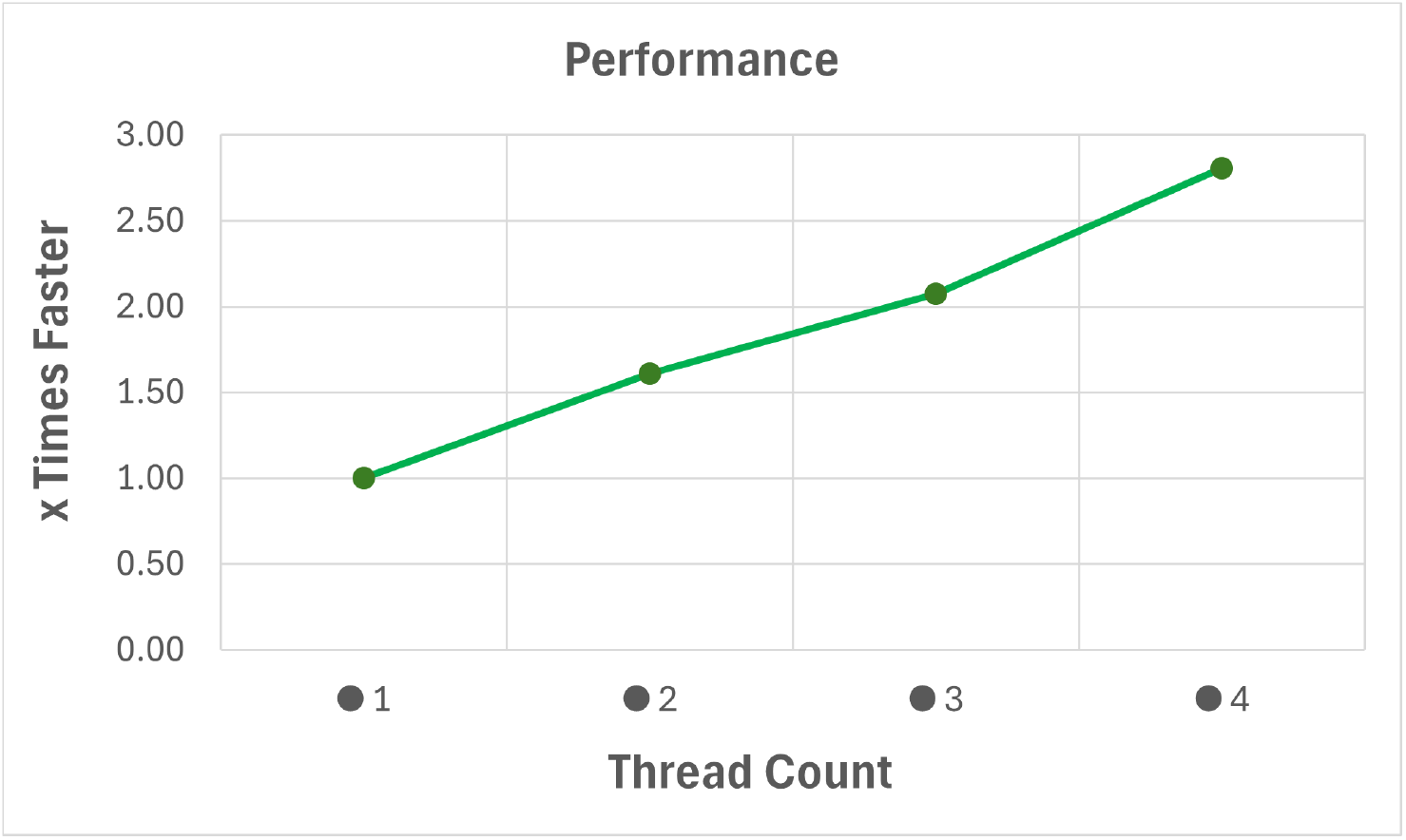
Performance scaling of Medlib under increasing thread counts. Execution time decreases as the number of threads increases, achieving near-linear speedup across 1 to 4 threads. This demonstrates the efficiency of Medlib’s thread-level parallelism, with consistent results across all configurations.

Using a single thread, the total alignment task requires 460.78 ms (2.91 μs per alignment). Doubling the threads reduces execution time to 285.84 ms, achieving a 1.61× speedup. With 3 and 4 threads, runtime further improves to 222.02 ms and 164.26 ms, corresponding to 2.08× and 2.81× performance increases, respectively, relative to single-threaded execution.

Importantly, the total number of alignments (158,428) remains constant across all thread counts, confirming that Medlib’s parallel implementation is deterministic and does not compromise correctness or result reproducibility. These findings highlight the utility of Medlib’s multi-threaded architecture for accelerating large-scale workloads without altering alignment precision.

## Discussion and conclusion

In this study, we introduce Medlib, a highly configurable and efficient C/C++ library for exact pairwise sequence alignment, designed to extend the capabilities of existing tools such as Edlib and Parasail. Built upon the foundation of Myers’s bit-vector algorithm, Medlib combines advanced compile-time dispatching, multi-modal input support, and customizable scoring with robust multithreading and novel alignment modes—providing both performance and flexibility for diverse bioinformatics applications.

Our benchmarking results demonstrate that Medlib consistently outperforms Edlib, its closest functional counterpart, across multiple alignment scenarios. In particular, Medlib’s Minimum mode executes 1.64× faster than Edlib while yielding over 6 million additional valid alignments, suggesting that Edlib may fail to capture certain optimal results under practical workloads. Furthermore, Medlib introduces a Threshold mode—currently unsupported by any other known exact alignment tool—that retrieves all alignments within a user-defined edit distance. Remarkably, this mode not only increases alignment yield by more than twofold compared to Edlib but also executes in less time, underscoring the effectiveness of Medlib’s internal optimisations.

Additional features such as overlap-aware reporting and motif boundary sensitivity offer users greater interpretability and control over result granularity. Our analysis shows that Medlib’s Distinct mode filters overlapping alignments to reduce redundancy with minimal performance cost, while motif-aware alignment options enable biologically informed trimming of flanking mismatches. These tools are particularly relevant in contexts like barcode demultiplexing, motif annotation, or amplicon detection, where precise boundary delineation is essential for downstream analyses.

Medlib also scales effectively with increasing computational resources. Our multithreading benchmarks reveal near-linear speedups up to four threads, reducing execution time by nearly threefold without compromising result determinism. This is achieved via dedicated threading strategies optimised for each alignment type, enabling Medlib to handle high-throughput workloads efficiently.

Despite these advances, certain limitations remain. Currently, Medlib supports only single-gapped affine models with scalar scoring; while sufficient for many DNA-based analyses, more complex substitution matrices or probabilistic scoring models (e.g., HMMs) are not yet integrated. These limitations are deliberate trade-offs aimed at preserving the library’s performance profile and will be addressed in future work.

Overall, Medlib provides a significant leap forward in exact sequence alignment by combining high performance with unmatched flexibility. Its unique features—such as threshold-based alignment, configurable boundary treatment, and deterministic multithreading—position it as a robust, general-purpose alignment library suitable for integration into modern bioinformatics pipelines. Medlib is freely available and open-source, facilitating widespread adoption and future extension by the research community.

## Code availability

Medlib is implemented in C/C++ programming language and is available at https://github.com/Mehdi-Rafeie/Medlib.

## Authors’ contributions

MR performed algorithmic analysis, code development, and writing – original draft. ORF provided biological insights, supervision, writing – review & editing. FV provided computational insight, review and editing.

## Acknowledgements

This work was supported by grants from Cancer Council NSW (RG 20-11) and UNSW Cellular Genomics Futures Institute.

## Supplementary material

Supplementary material associated with this article can be found in the attachment.

