## Supplementary data for "Medlib: A Feature-Rich C/C++ Library for Exact Alignment of Nanopore Sequences Using Multiple Edit Distance"

#### 1. Alignment Types

Table S1 summarizes the 15 alignment types supported by Medlib, defined by all valid combinations of query and target input modalities. Each alignment type is implemented as a distinct module, facilitating independent compilation and integration into diverse analysis pipelines.

Table S1. Alignment Types Supported in Medlib.

| **Alignment Type** | **Description** |
|---|---|
| SQST | Single Query to Single Target |
| SQMT | Single Query to Multiple Targets |
| SQTFtxt | Single Query to Target File (Text) |
| SQTFfa | Single Query to Target File (FASTA) |
| SQTFfq | Single Query to Target File (FASTQ) |
| MQST | Multiple Queries to Single Target |
| MQMT | Multiple Queries to Multiple Targets |
| MQTFtxt | Multiple Queries to Target File (Text) |
| MQTFfa | Multiple Queries to Target File (FASTA) |
| MQTFfq | Multiple Queries to Target File (FASTQ) |
| QFST | Query File to Single Target |
| QFMT | Query File to Multiple Targets |
| QFTFtxt | Query File to Target File (Text) |
| QFTFfa | Query File to Target File (FASTA) |
| QFTFfq | Query File to Target File (FASTQ) |

#### 2. Motif Boundary Treatment Examples

In many alignment tasks involving barcodes or other short motifs, mismatches at the motif boundaries may arise due to sequencing error, truncation, or imperfect primer annealing. These mismatches can lead to biologically meaningful bases being incorrectly excluded from the alignment if treated as strict substitutions.

In the left panel of Figure S1, an ordinary motif is aligned with a target sequence using default settings (No Flanking mode), where the leading C is treated as a mismatch and excluded from the alignment. In the middle panel, this region is identified as a flanking motif adjacent to a barcode. Under default alignment, the mismatch still prevents the barcode from extending to its full length. The right panel shows the effect of enabling Flanking Start mode. In this mode, leading mismatches at the motif boundary are reinterpreted as deletions from the upstream sequence rather than penalized substitutions. As a result, the initial C can now be included in the alignment, correctly extending the barcode boundary. This flexible reinterpretation improves alignment sensitivity in scenarios where barcodes or UMIs are adjacent to engineered constant regions or primers, reducing the risk of truncation due to marginal mismatches.

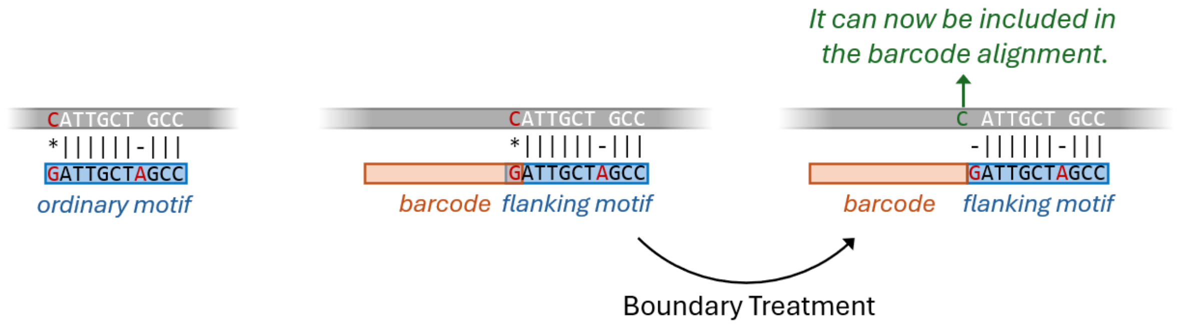


**Figure S1.** Illustration of boundary treatment in Medlib for barcode-containing motifs.

### 3. Multi-Threading Strategies

Table S2 provides an overview of the multi-threading strategies implemented for each Medlib alignment type. Medlib optimizes parallelism by tailoring its threading models to the specific data access patterns and computational demands of different input modalities.

**Table S2.** Multi-threading strategies tailored to each Medlib alignment type.

| Alignment Type | Multi-Threading Strategy |
|---|---|
| SQST | N/A |
| SQMT | Target Parallel Partitioning |
| SQTFtxt | Target Producer-Consumer Model |
| SQTFfa | Target Producer-Consumer Model |
| SQTFfq | Target Producer-Consumer Model |
| MQST | Query Parallel Partitioning |
| MQMT | Sequential Query, Target Parallel Partitioning |
| MQTFtxt | Target Producer-Consumer Model |
| MQTFfa | Target Producer-Consumer Model |
| MQTFfq | Target Producer-Consumer Model |
| QFST | Query Producer-Consumer Model |
| QFMT | Sequential Query, Target Parallel Partitioning |
| QFTFtxt | Sequential Query, Target Producer-Consumer Model |
| QFTFfa | Sequential Query, Target Producer-Consumer Model |
| QFTFfq | Sequential Query, Target Producer-Consumer Model |

In this table, "Target Parallel Partitioning" and "Query Parallel Partitioning" involve static division of targets or queries among available threads, minimizing synchronization overhead. "Target Producer-Consumer Model" and "Query Producer-Consumer Model" refer to multi-threading schemes in which one or more producer threads sequentially read targets or queries from memory or files, while multiple consumer threads simultaneously perform alignments on the incoming data. This decouples I/O and computation, enabling continuous data feeding to CPU cores without bottlenecks and significantly improving throughput, especially for large files. "Sequential Query" indicates that queries are processed serially to ensure correctness when parallelizing over targets.

### 4. Medlib Configuration

Figure S2 displays an example of the configuration tree of Medlib.

```
Medlib Configuration
│
├──Input/Output Config
│   ├──Query Input Type: Multiple
│   ├──Target Input Type: FASTQ File
│   ├──Input
│   │   └──MQTF
│   │       ├──Queries: Forward and reverse regions (flanking and constant)
│   │       └──Query Count: 14
│   └──Output
│       └──Store in Memory: No
│
├──Alphabet Config
│   └──Alphabet Name: ACGT [Medlib Predefined Alphabet]
│
├──Operation Config
│   ├──Align Mode: Local
│   ├──Operation Mode: Threshold
│   ├──Objective: Range
│   └──Custom Operation
│       └──UDF Type: Batch UDF
│
├──Edit Distance Config
│   ├──Edit Distance Type: Collection
│   └──Edit Distance: 10% of the query length
│
├──Motif Config
│   ├──Overlap Handling: Distinct
│   ├──Motif Treatment Type: Specific for Each Motif
│   └──Motif Treatment:
│       ├──FF1 = Not Flanking
│       ├──FC1 = Flanking →¦
│       ├──FC2 = ¦← Flanking →¦
│       ├──FC3 = ¦← Flanking
│       ├──FC4 = ¦← Flanking →¦
│       ├──FC5 = ¦← Flanking
│       ├──FF5 = Not Flanking
│       ├──RF5 = Not Flanking
│       ├──RC5 = Flanking →¦
```

```
│              ├──RC4 = ¦← Flanking →¦
│              ├──RC3 = Flanking →¦
│              ├──RC2 = ¦← Flanking →¦
│              ├──RC1 = ¦← Flanking
│              └──RF1 = Not Flanking
│
└──Parallelism Config
   └──Threading: Multi-Threaded
```

**Figure S2.** Medlib Configuration Example
